# Immersive display systems for simulating natural vision

**DOI:** 10.64898/2026.07.27.741046

**Authors:** Harish Katti, Aidan P. Murphy, Michael Helde, Harshawardhan Deshpande, Tyler J. Lee, Rekeya Knight, Carly Gregg, Cristina Solinas, Katherine Cameron, Daryl Bandy, George Dold, David A. Leopold

## Abstract

Vision is traditionally studied using simplified stimuli presented briefly near the center of a flat display while subjects fixate. This paradigm contrasts sharply with real-world vision, which is immersive, dynamic and strongly entrained to self-initiated actions. Traditional approaches have enabled systematic study of the neural encoding of visual features and the influence of cognitive operations such as attention. However, other methods are needed to study more holistic and first-person perspectives on vision, such as those related to physical space, continuous time, and self-movement. To enable the study of these aspects of real-world vision, we developed hemispherical (“dome”) display systems for macaque visual neuroscience. For functional MRI experiments, a compact rear-projection dome display fits within the bore of the scanner. For electrophysiology, a larger front-projection dome display is illuminated from above using a spherical mirror. Both systems provide stimulation of approximately 180° of the subject’s field of view, enabling visual immersion. To ensure accurate angular geometry across the hemispherical display, we present unified rendering and calibration procedures that support natural fisheye videos, conventional visual stimuli, and virtual 3D environments. The calibration procedure automatically compensates for projector, mirror, and dome distortions through geometric pre-warping, ensuring correct visual-angle representation across the display. Pilot fMRI experiments demonstrate robust activation of peripheral visual cortex during both conventional pattern stimulation and naturalistic self-movement. Together, these dome systems provide a flexible platform for investigating aspects of vision that are difficult to study with conventional displays, including peripheral processing, visual immersion, optic flow, self-motion, and holistic scene perception.

## 1. Introduction

Our visual experience of the world is immersive: we exist within and move through a three-dimensional (3D) environment, at each moment facing a visual panorama that occupies our entire field of view. Amid this immersive experience of the world, natural locomotor and reorienting behaviors create visual events that dynamically stimulate the entire retina. The brain must discount some aspects of this visual stimulation and utilize other aspects (Kim et al., 2016; Wurtz, 2008). The core operations associated with immersion and self-generated actions are deeply rooted in evolution and broadly shared by animals that use vision to guide behavior (Leopold et al., 2020). The neural mechanisms underlying these fundamental aspects of vision are not well studied, in part because traditional stimuli and paradigms typically focus on central feature representation. However, specialized display systems utilizing geometrically valid full-field stimuli can compellingly convey the experience of inhabiting and navigating a 3D environment (Shapcott et al., 2025a). An immersive, full field display allows for the systematic study of peripheral vision (Figure 1A-C), along with the spatial and temporal composition of a scene viewed from a geometrically valid first-person perspective (Figure 1D-F). This combination opens new doors studying how the brain supports fundamental but minimally explored aspects of visual cognition.

**Figure 1.**
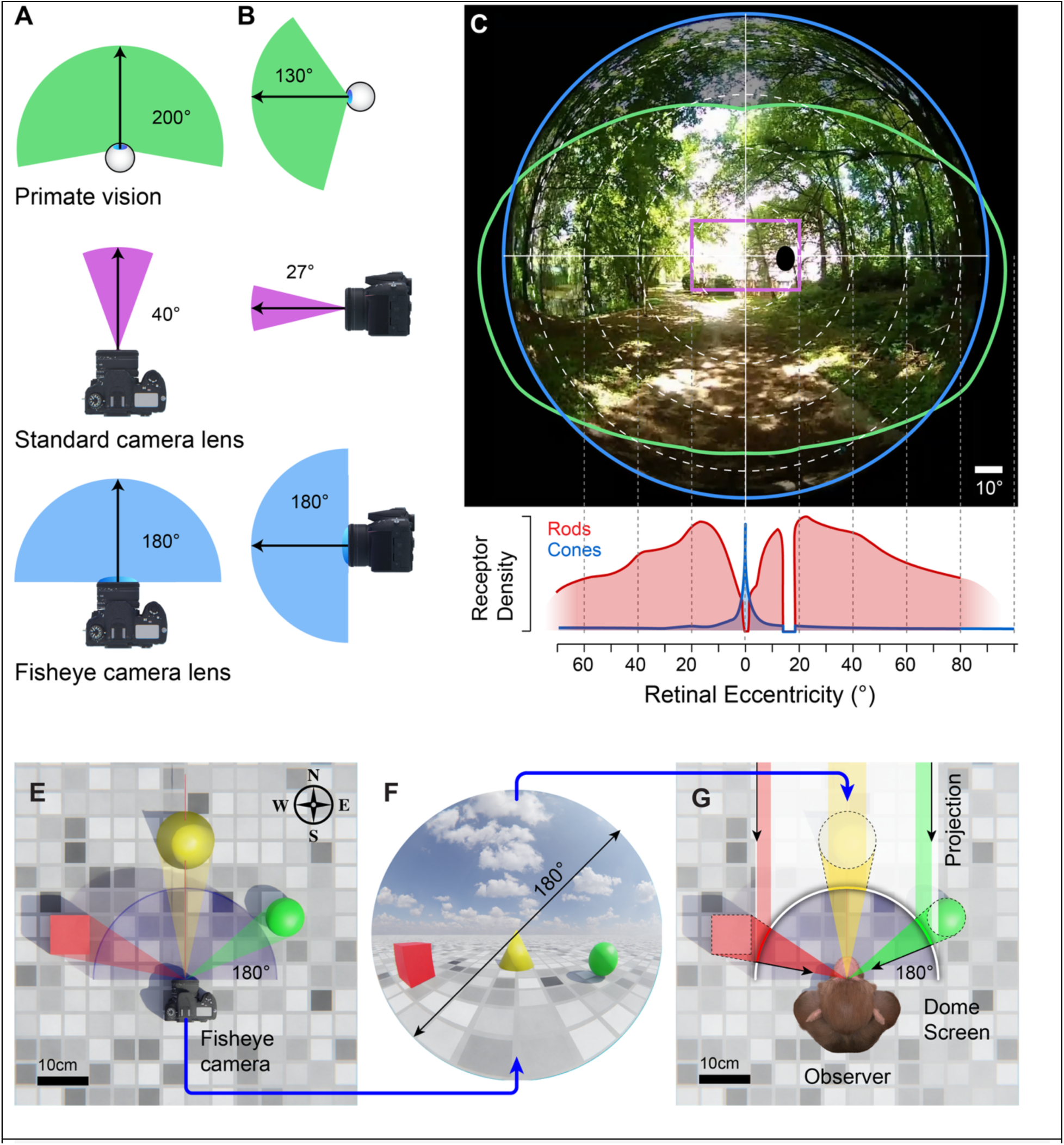
Comparison of visual field coverage and preservation of geometric relations. A – C. The primate visual field (green) extends far beyond the fovea, with an estimated horizontal span of 200° (A) and vertical span of 130° (B). In contrast, traditional camera lenses and sensors are designed to capture relatively smaller fields of view. A standard camera lens with 50mm focal length (magenta) covers approximately 40° horizontally and 27° vertically. Conventional image capture and display technologies are therefore ill suited to providing naturalistic visual stimulation of the entire visual field. Panoramic fisheye lenses (blue) can capture 180° (or more) of the visual field in both the horizontal and vertical dimensions and can thus provide visual stimulation that more closely approximates the experience of natural primate vision. E – G. By matching the angular field-of-view (FOV) of image generation or capture (E) to that of image display (G), natural real-world geometric relations –including metric object size (e.g. centimeters)—are preserved. E. A birds eye view of a virtual scene containing a fisheye camera, which captures a 180° field-of-view image (F). When this image is correctly projected onto a 180° hemispheric dome screen, the spatial scale and relations of the original environment are preserved, and an observer located at the center of the dome display perceives the scene in a first-person manner, as though they were physically present in it.

Experimentally, immersive stimulation requires a visual display system that permits electrophysiology or fMRI measurements using specially constructed full-field scenes, dynamic videos, or virtual environments. Electrophysiological studies in primates examining large portions of the visual field have typically used flat screens (Vaziri et al., 2014a; Wild & Treue, 2021). Studies in other species have employed arrays of multiple flat-screen displays (Ogawa et al., 2025), or curved projection screens (Hölscher et al., 2005; Katayama et al., 2012). For humans, head-mounted displays (HMDs) provide increased visual field coverage over traditional displays (Xiao et al., 2025a). These commercial units cannot easily be tailored to the smaller anatomical proportions of monkeys, nor are they compatible with the requirements of fMRI or electrophysiology recordings. However, several groups have devised other approaches for wide-field visual stimuli for human fMRI (Greco et al., 2016; Jolly et al., 2021; Mikellidou et al., 2017; J. Park et al., 2024). For full immersion, a particularly attractive approach is the projection of full-field scenes onto a hemispheric ‘dome’ surface. This method has a long history of use in planetariums (Chant, 1935a), and has previously been adapted for smaller scale settings (P. Bourke, 2005a). Several groups have used visual projection onto hemispheric domes for primate electrophysiology research (Shapcott et al., 2025b; Yu et al., 2015). In this article, we provide detailed descriptions of two hemispheric dome display systems we developed for macaque neuroscience research. The first system consists of a compact (46 cm diameter) back-projected dome that fits inside an MR scanner and is used for functional MRI. The second system is a larger (193 cm diameter), front-projected hemispheric dome and is designed for electrophysiology experiments. We report the design and construction of these setups, along with details about stimulus creation and calibration procedures to facilitate geometrically accurate real-world simulation. We survey the challenges and opportunities presented by this testing paradigm.

## 2. Methods

Traditional visual display technologies (e.g. flat 2D screens) cannot simulate the immersive nature of natural vision. Modern commercially available head-mounted display (HMD) technologies have come closer to providing an immersive visual experience and allow for free movement. However, head-mounted devices are designed specifically for adult human anatomical proportions and would be challenging to adapt to neuroscience studies in nonhuman primates. We therefore sought to construct hemispheric ‘dome’ screen projection systems for two different neural measurement modalities that impose differing restrictions: functional magnetic resonance imaging (fMRI) and extracellular electrophysiology.

### 2.1. Hardware configurations

#### 2.1.1. A rear-projection dome for fMRI

##### 2.1.1.1. Motivation

To examine fluctuations in brain activity during naturalistic immersive vision, we sought to present full-field visual stimuli to awake macaque subjects in an MRI scanner (Siemens Prisma). The primary constraints of the MR-environment are the high magnetic field strength (in this case 3 Tesla) and the spatially restrictive magnet bore (in this case 60cm diameter). The traditional solution to these constraints is to project video content from a projector located away from the magnet onto the rear surface of a small flat screen that can fit inside the magnet bore. Human subjects typically view the projection screen through a mirror while lying in a supine position (although see Jolly et al., 2021; Park et al., 2024), while awake macaque subjects typically view the screen directly with their head upright in a ‘sphinx’ position. We adapted this standard approach to the use of a hemispheric dome screen as described below (Figure 2A-D).

**Figure 2.**
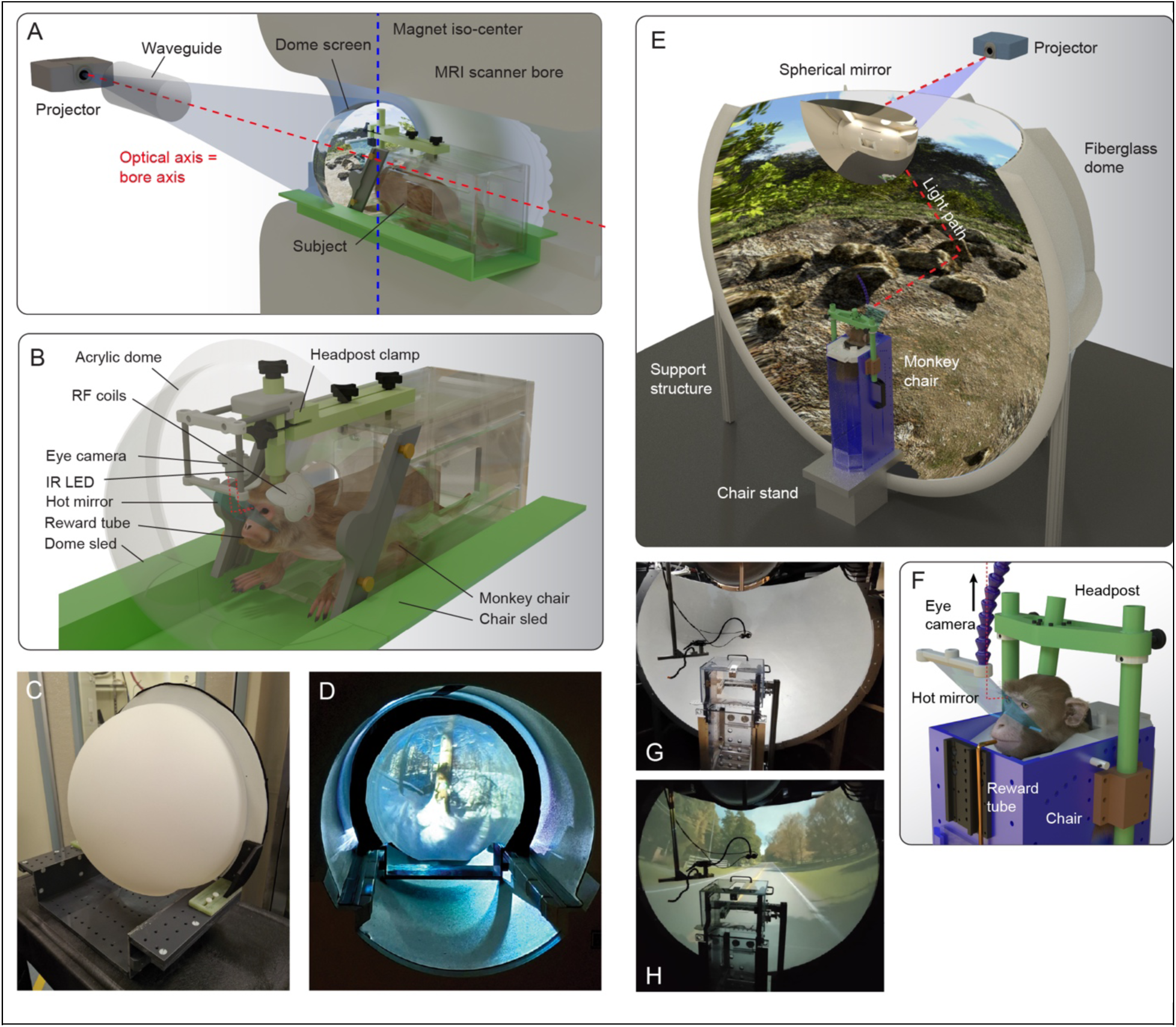
Illustrations of the hardware setups for (A-D) rear-projection dome screen for fMRI and (E-H) front-projection dome screen for electrophysiology. A. The rear-projection dome screen for fMRI is inserted inside the magnet bore, so that it surrounds the animal’s head, which is fixed at the magnet’s isocenter. B. Illustration of a macaque subject sat in a sphinx position for fMRI studies, with necessary hardware (headpost restraint, RF coils, video eye tracking and reward delivery systems) fitting within the concave volume of the dome screen. C. The rear side of rear-projection dome, outside the bore. D. The viewing side of the rear-projection dome inside the bore. E. The front-projection dome screen for electrophysiology, with the light path (red dashed line) indicated from projector to dome surface via ‘spherical’ mirror. F. Video eye tracking is achieved via infrared light reflecting off a hot mirror held at 45° in front of the animal’s face, allowing the camera (not shown) to be positioned away from the recording electrodes without obstructing the visual field. G. Photograph of the front-projection dome setup and (H) with a scene projected onto it.

Several spatial constraints are imposed by the combined needs for high quality brain imaging and full-field visual stimulation. For optimal imaging of the brain, the subject’s head must be positioned at the magnet’s isocenter, which is located deep inside the magnet bore. For a hemispheric dome screen to produce an immersive ∼180° field of view, the outer most edges must be positioned level with the observer’s eyes (i.e. 90° eccentricity). We therefore selected a hemispheric dome that was small enough to be inserted deep inside the magnet bore, but large enough to accommodate the subject’s head and associated experimental apparatus.

##### 2.1.1.2. Hardware

###### Acrylic hemispheric dome

A clear, free-blown acrylic hemispheric dome measuring 45.7cm (18”) in diameter, with a 3.8cm (1.5”) flange, was purchased (CDS, Laird Plastics, CA). The outer diameter of the hemisphere was 53.3cm, thus mostly filling but still easily clearing the 60cm diameter bore of the Siemens Prisma 3T MRI scanner. The fabrication method for this product starts with a uniform 0.25” thick sheet of clear acrylic that is heated to over 220°C before air is blown inside to stretch the plastic to shape. This approach results in high optical clarity, but gradual reduction in thickness across the surface from the outer edge (∼3/16”) to the center (∼1/16”). To stand the dome screen vertically upright inside the scanner bore, a custom ‘sled’ was constructed from Garolite (G10) fiberglass panels and low-friction Delrin guides, which sat on and slid along the scanner’s patient bed rails inside the bore (Figure 2C). Two corners were waterjet cut from the acrylic hemisphere to fit the sled and holes were drilled around the flange to create mounting points for securing the dome inside the scanner bore. Custom designed brackets were 3D-printed and attached to the sled and the flange of the dome via nylon ¼”-20 machine screws. Finally, the rear (convex) surface of the dome was sprayed with rear projection paint (Screen Goo, Goo Systems Global, Canada), creating a uniform translucent matte white surface, while the inner (concave) surface was left untreated.

###### Projector system

A projector (Epson PowerLite Pro G5450WU, 1080 x 1920 pixels, 60Hz refresh rate) fitted with an after-market zoom lens (Navitar PT-D5500, 6”-9” zoom; D:W 10.7 - 16; focal length 150 - 230mm) located outside of the magnet room was projected through a 20 cm diameter waveguide into the magnet room. Due to the location of the equipment room, the light path from the projector began orthogonal to the magnet bore and was reflected off a large (60 cm x 40 cm) front-surface mirror (Edmund Optics, NJ, USA) mounted to the wall at a 45° angle and located at the end of the magnet bore. The total projection distance from projector to screen (via mirror) was 420 cm. The optical axis of the projector was level with the center of the waveguide and the center of the magnet bore (Figure 2A). The projector was mounted on a mechanical height-adjustable frame (UpLift Desks, TX, USA), with a custom extruded aluminum frame (8020 Inc.) allowing adjustments of projector pitch and yaw angles. Spatial calibration of the projector was checked and adjusted before each experiment by projecting an image of a polar grid with central crosshairs onto the dome surface and manually assessing alignment.

Note that as projection distance increases, the projection frustum approaches an approximation of orthographic projection. This minimizes the surface area of the dome screen that has a surface normals tangential to the light path at the far peripheral edges. However, it is impossible to obtain a complete 180° visibility of the projected image on the dome surface using this single rear projector method, since some portion of the dome surface will necessarily be oriented orthogonal to the light path.

###### Eye tracking

When projecting visual stimuli onto a traditional flat rear-projection screen, video eye tracking can be accomplished by placing a camera fitted with a telephoto lens outside of the magnet bore. However, this approach requires a hole somewhere in the screen for infrared light to pass from the illuminator to the subject’s eye and back to the eye camera. Since the additional of a hole in the dome screen would not only diminish visual field coverage of stimulation but also disrupt the subject’s sense of immersion, we instead opted for a different approach. An MR-compatible video camera (MRC Systems GmbH, Germany – model 12M) and infrared LED were positioned above a 45° hot mirror (Edmund Optics, NJ, USA), with a contour waterjet cut along one edge to accommodate placement over an adult macaque’s nose (Figure 2B). All parts were held in place by custom-designed 3D-printed plastic hardware, attached to the headpost holder via modular hose (Loc-Line, OR, USA). Data from the video camera passed via an RF filter panel to a PC located in the equipment room, running Arrington ViewPoint video eye tracking software.

#### 2.1.2. Front-projection dome for electrophysiology

##### 2.1.2.1. Motivation

Since commercial HMDs designed for humans cannot be easily adapted to the smaller inter-pupillary distances of non-human primates, we sought alternative approaches to wide field visual stimulation. The concept of front-projection onto a hemispheric dome screen has been used in planetariums for over a century (Chant, 1935b). A more recent development of hemispheric front-projection was developed by Bourke (2005), which removes the expense and technical hurdles associated with either panoramic fisheye projection lenses or multi-projector setups. Instead, a relatively lower cost ‘spherical’ mirror is used to distribute the projected image from a standard commercial projector onto the inner surface of a vertically oriented hemispheric projection dome.

We modified the original ‘iDome’ design (P. Bourke, 2005b) in several ways to better suit our specific experimental needs and available laboratory environment. First, we scaled the dimensions of the dome down slightly (from the original 3m to 2m in diameter) to fit inside our existing Faraday shielded electrophysiology recording booth. Second, we decided to invert the dome, so that the truncated edge, mirror and projector would be located near the ceiling rather than on the floor. This decision was driven primarily by the need to protect the delicate front-surface mirror from dirt and splashes of liquid reward given to the animals during experiments. However, this approach also has the benefit of providing greater coverage of the lower visual field, which is more extensive than the upper visual field for a head-fixed macaque due to occlusion by the prominent supraorbital margin. Finally, we decided not to add a circular hole for the projector at the truncated rim of the dome, which would have complicated fabrication. In the original non-inverted iDome design, this hole is necessary because the truncated edge of the dome rests on the ground, leaving nowhere for the light to pass through. In our inverted design, the projector is located above the truncated ridge and the projector’s vertical lens shift is used to adjust the light path downwards to the mirror.

##### 2.1.2.2. Hardware

We ordered a 2-meter (6’4”) diameter hemispheric fiberglass (Model #A-139, Architectural Fiberglass, Inc., OH), which shipped in 8 segments and required assembly and finishing. The four segments that made up the upper half of the dome were truncated at the line corresponding to +45° elevation, per our customization request. On arrival, the segments were bolted together (3/8” x 2” grade 5 bolts) and mounted on extruded aluminum profile legs. The interior surface was caulked, plastered and painted with silver paint developed specifically for polarized 3D projection (Screen Goo, Goo Systems Global, Canada). The finishing and painting required specialist equipment (for paint spraying and proper ventilation, etc.), and we therefore outsourced this step to a professional with experience in automotive body paint.

We ordered a 60cm diameter first surface half dome projection mirror (Acril Convex Mirrors, Australia; although a similar product is available domestically in the US from Go-Dome^TM^, Avela Corp., TX). The mirror surface is coated in a thin polymer for limited protection, and the mirror should not be touched and cleaned only with air. Our decision to invert the original iDome design and suspend the mirror *above* the subject rather than below, was partially motivated by the need to preserve the optical clarity of this critical element of the system.

There are several criteria to consider when selecting a suitable projector for iDome projection. The primary requirement is the interaction of projector focal length and available space for positioning the projector such that it can form a ∼40 to 50cm wide image on the 60cm diameter front surface mirror. We additionally wanted a projector with a minimum of 1920 x 1080 pixels native resolution at 60Hz, and the capability of presenting stereoscopic 3D using passive polarizing filters. We found the Optoma UHZ50 DLP projector to fit all these criteria since it has a VESA sync port for synchronizing stereoscopic 3D hardware and can display 4K images at 60Hz (or 1080p images at 240Hz).

### 2.2. Visual stimulus generation

Images and videos taken with a conventional camera do not capture sufficient angular subtense of the visual scene to mimic immersion, as their field of view is limited compared to that of the retina (Figure 1). While such images can be scaled to cover the entire retina, this does not faithfully convey the natural angular geometry of the scene. For this, one needs to acquire panoramic images using a wide-field (fisheye) lens that matches the angular subtense of the display (Figure 1E-G). In this way,

Any point in a fisheye image corresponds to a unique three-dimensional viewing ray, represented by its azimuth and elevation on the viewing sphere. For real fisheye cameras (Figure 3A-B), the mapping between image pixels and viewing directions depends on the lens projection function (e.g., orthographic) and residual lens distortions, which are estimated through camera calibration to establish an accurate pixel-to-ray correspondence. In contrast, fisheye images generated in virtual environments (Figure 3C-D) are synthesized analytically from the underlying 3D scene geometry and therefore require no lens calibration. Instead, the desired fisheye projection model is explicitly defined during rendering, ensuring that every pixel corresponds to a known viewing direction. In both cases, the resulting fisheye image provides a one-to-one mapping between image pixels and angular viewing directions, making it a natural representation for hemispherical dome projection and angularly accurate stimulus generation.

**Figure 3.**
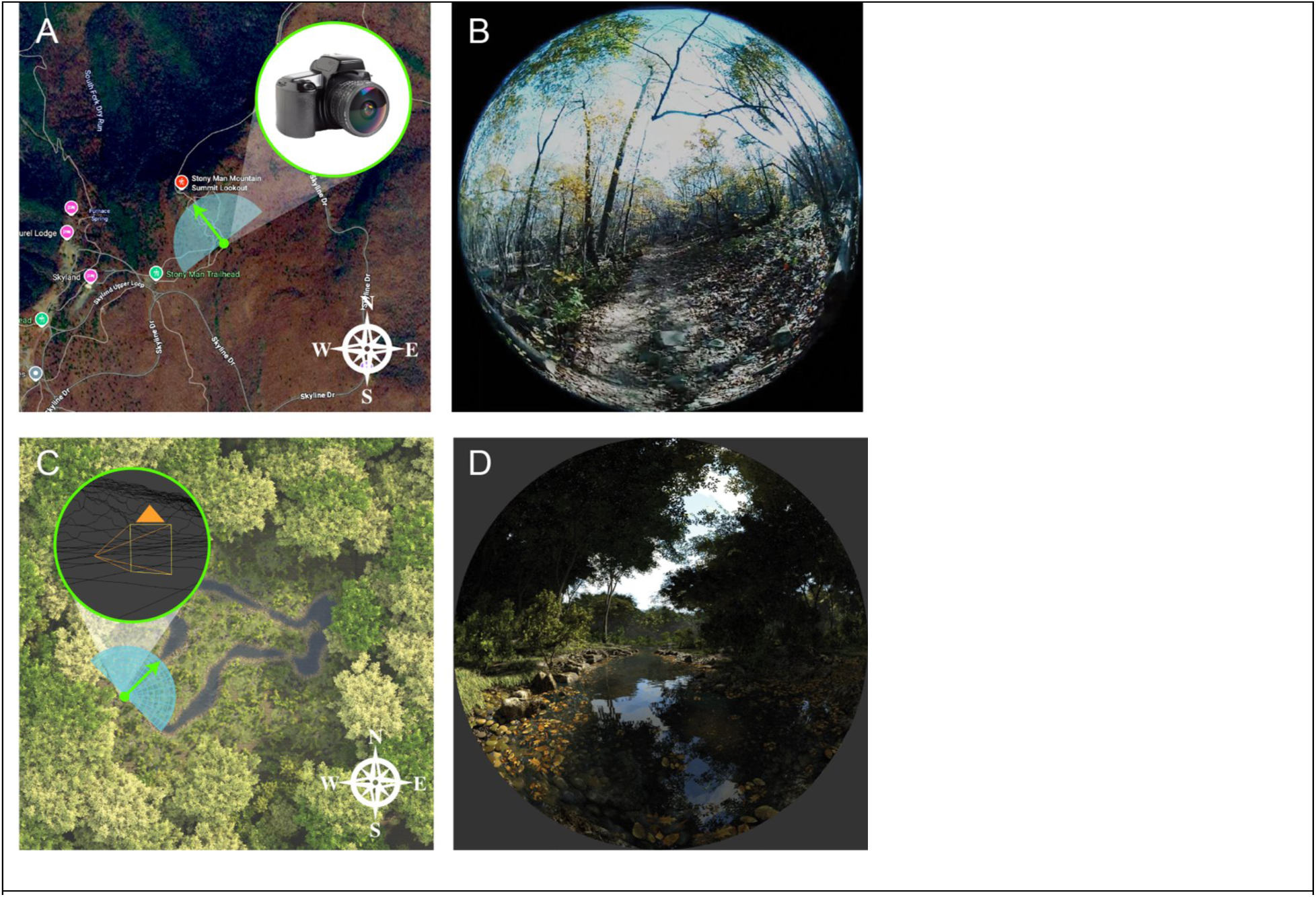
Capturing 180° fisheye views in real and virtual environments. (A) A birds eye view of a real-world location where a hardware fisheye camera was used to capture a 180° fisheye image (B) at a given allocentric location and compass direction (green arrow in A). (C) A bird’s eye view of a virtual forest environment in Blender, illustrating the allocentric location of a virtual camera and it’s direction (green arrow).

#### 2.2.1. Fisheye acquisition with camera

Affordable and compact consumer video cameras with ultra-wide angle (fisheye) lenses have become more popular for virtual reality (VR) content creation. We used a commercial stereoscopic digital camera (Lucid Cam) with fisheye lenses of 1.9mm fixed focal length, capable of recording video at 2160 x 2160 pixels (per eye) and 30 frames per second. Camera motion was partially stabilized during video recording by mounting the camera on a 3-axis handheld gimbal (Zhiyun SmoothQ) via a custom 3D-printed bracket.

#### 2.2.2. Virtual Reality

There are numerous software options for generating and animating VR environments. These can either be ‘real-time’ game engines, such as Unity (P. D. Bourke, 2005) or Unreal Engine (Shapcott et al., 2025b) that allow for user-interaction or closed-loop experiments, or off-line render engines, such as Blender (P. D. Bourke & Felinto, 2010) that provide more fine-grained controllable parameters and more photorealistic renders. Virtual scenes are typically composed of surface mesh geometry with materials and shaders that determine object appearance, lighting and a virtual camera that defines the perspective of the rendered output. The location, movement, and other parameters of virtual cameras can be finely controlled, including the ability to produce 180° fisheye rendered outputs suitable for hemispheric dome display.

##### 2.2.2.1. Offline VR Rendering

Offline rendering of VR scenes allows more raytracing computation time per frame, resulting in higher quality and more photorealistic renders. Blender (www.blender.org) is a powerful and well established open-source 3D animation software, that is widely used, including in previous neuroscientific research (Khandhadia et al., 2021, 2023a; Murphy & Leopold, 2019) and dome projection settings (P. D. Bourke & Felinto, 2010). It is controllable via both the graphical user interface and Python commands (Blender installs with its own Python interpreter), which simplifies parametric control of experimental stimulus variables. To render images of virtual 3D scenes suitable for hemispheric dome projection using Blender (version 5.0), various settings of the virtual camera can be configured. Here we describe the steps to configure the camera using the GUI, and below we provide a code snippet to achieve the same actions programmatically via Python.

For 180° fish-eye renders, the ‘Cycles’ render engine must first be selected. With a camera object selected, in the camera ‘Data’ tab of the Properties menu, the lens type should be set to ‘panoramic’ and ‘panorama type’ should be set to ‘fisheye equidistant’, which then allows adjustment of ‘field of view’ (°). The aspect ratio of the output frame images should be set at 1:1, with the pixel values matching (or exceeding) the vertical resolution of the projector. The rendered output will appear as a fisheye image within a circular aperture and black pixels in the corners of the image. These settings can also be configured via Python code as seen in the demo script *FisheyeSetup.py* provided in the supplementary materials.

##### 2.2.2.2. Real-time VR rendering

‘Game engines’ are video rendering software capable of updating displays of 3D virtual environments in real-time (i.e. at the refresh rate of the display). Real-time rendering opens a wealth of additional experimental approaches, including user interaction and neurofeedback, at the cost of slightly less realistic rendered output compared to offline renders. Most real-time engines can also be used to render animations off-line but with much faster turnaround than typical offline render engines.

Our approach for rendering fisheye images for dome projection in one such real-time 3D engine (Unity) uses a cubemap-based approach (P. Bourke, 2009). A dedicated World Camera captures the scene each frame by rendering a full cubemap (six orthogonal views covering 360°). The cubemap is then passed to a custom shader via an image-effect script attached to the Projection Camera. The shader remaps the cubemap into a circular fisheye image by converting screen-space pixels into angular coordinates and sampling the corresponding direction in the cubemap. Importantly, the image effect overrides all prior rendering, ensuring that the final output is entirely derived from the cubemap rather than the standard camera pipeline.

A separate controller component manages the parameters governing this transformation, including fisheye field of view (up to ∼270°), cubemap resolution, anti-aliasing, and fade functions that modulate intensity across the dome. The controller also enforces the orientation of the World Camera (via pitch and roll), ensuring that the cubemap aligns correctly with the fisheye projection. This architecture produces a geometrically consistent wide-field representation suitable for dome or immersive display systems, while maintaining flexibility in resolution and rendering quality through configurable cubemap sampling and post-processing. Examples of such fish-eye image generation is shown in Supplementary Figure S1.

As designing stimuli and experiments in the dome systems can employ a variety of software options that can have overlapping capabilities, we have organized and compared their capabilities in Table S1, based on our experience in using them for experiments in dome systems.

### 2.3. Image calibration for front-projection dome

The iDome design is an elegant solution to the problem of front-surface 180° fisheye projection (P. D. Bourke, 2005). Specifically, it solves the problem of both the observer and the projector needing to be co-located at the optical center of the dome by using a spherical mirror which is vertically offset from the center. However, this also introduces additional image distortion, which needs to be corrected in software before the image is sent to the projector (Figure 4).

**Figure 4.**
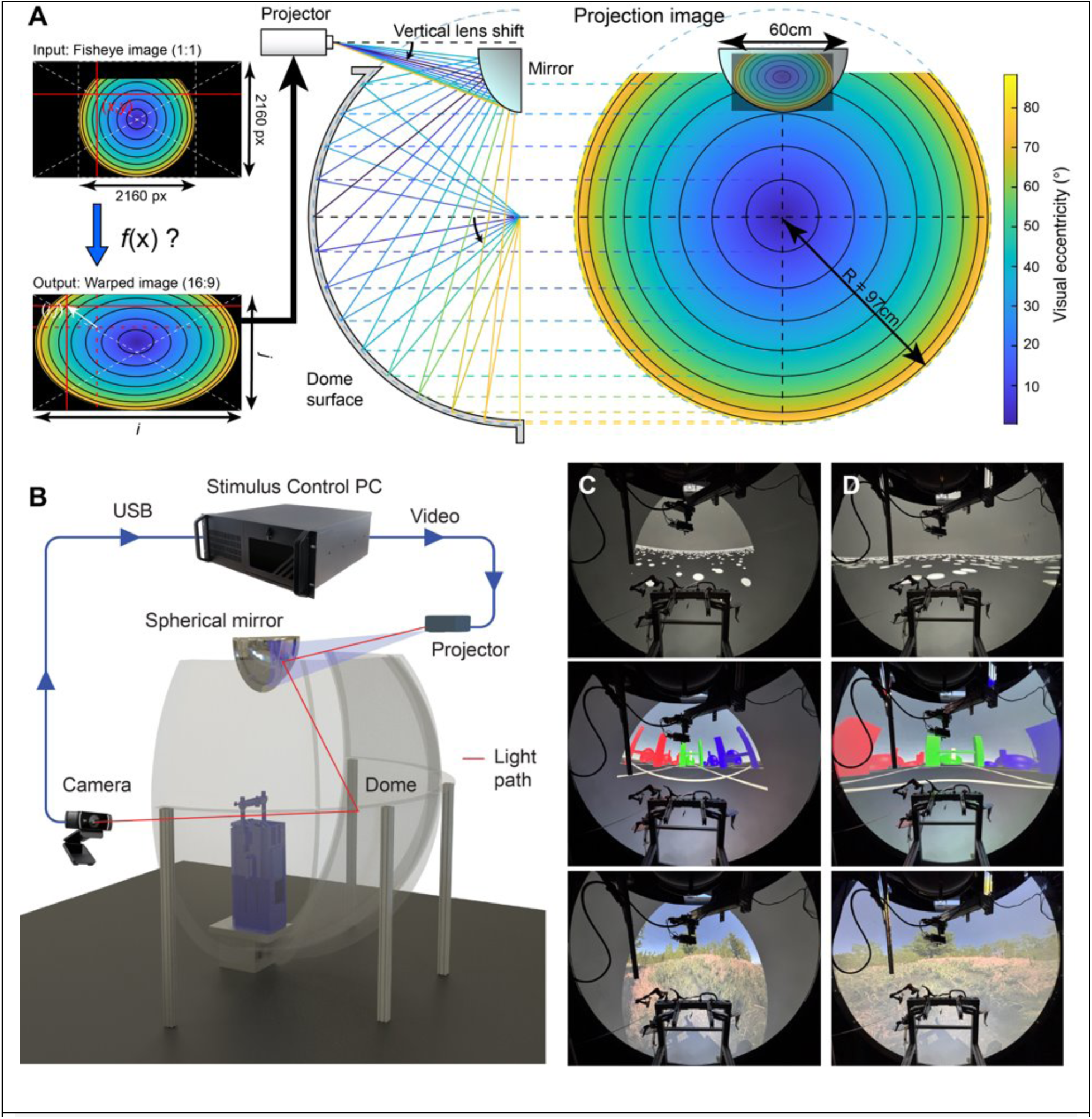
Mapping pixels to retinal angles for hemispheric dome projection. (A) A fisheye image (i) orthographically projected onto a hemispheric screen (iii) would preserve veridical geometry of the depicted 3D scene for an observer located at the center. However, the iDome system introduces multiple non-linear image distortions (iii). The goal of the pre-warping process is to derive the 2D mapping function (f(x)) that successfully preserves correspondence between the polar coordinates of the fisheye image and the dome surface projected image. (B) Schematic of the hemispherical dome projection and calibration setup. Visual stimuli generated on the stimulus control PC are sent to a projector, reflected off a spherical mirror, and projected onto the inner surface of the hemispherical dome. A USB camera positioned outside the dome is used for calibration and monitoring of the projected image geometry. (C, D) Elements such as the curved mirror produce non isotropic resampling particularly noticeable at large eccentricities, along the horizon, as shown in (C) and motivates the need for pre-warping and non-uniform resampling of immersive stimuli to recover correct 3D geometry (D).

The extent of angular shift for a projected pixel in the stimulus can be estimated manually by projecting a single dot of known angular subtense and comparing it against known angular positions on the dome that have either been marked physically (e.g.; using a mark or laser dot on the dome) or using a custom apparatus that we initially designed. Although this allowed us to estimate the shift of projected points, the procedure is tedious and time consuming as one needs to repeat the procedure for a relatively large number (121 in our case) of points that form the reference pattern. This manual process was also error prone at the far periphery and resulted in artifacts in the final displayed image. We therefore sought to automate the process using a computer vision approach.

#### 2.3.1 Computer vision-based closed loop automatic distortion estimation

Automated calibration was accomplished by through a closed-loop system consisting of a webcam monitoring the position of sequentially presented dots. First, following a geometric unwarping procedure, a polar calibration pattern consisting of concentric rings (5°–90° in 5° steps) and radial lines (0°–360° in 10° steps) was generated at the native projector resolution (1920 × 1080) and displayed using Psychtoolbox (Brainard, 1997; Pelli, n.d.). Then, for each ring–line intersection, a single red dot was rendered sequentially on the projector while a fixed webcam recorded the physical dome surface. We projected dots using Psychtoolbox routine *<u>DisplayUndistortionBVL</u>* for estimation of such distortion fields. The dot location in camera coordinates was extracted by background subtraction and maximum-intensity detection, yielding paired correspondences between desired screen coordinates and observed dome coordinates.

All dome coordinates were re-centered relative to the dome apex and rescaled to screen units using the measured dome radius in the camera image. These matched point sets were used to estimate both forward (screen to dome) and inverse (dome to screen) mappings. Mappings were learned using robust second-order polynomial regression (MATLAB fitlm, poly22 model). The inverse mapping was used to pre-warp stimulus images by remapping each pixel in the source image to its predicted screen location prior to projection. The resulting pre-warped images, when projected onto the dome, produced visually undistorted stimuli in angular coordinates.

#### 2.3.2. Eye tracking in the front projection dome

Conventional video eye-tracking for vision experiments uses a small angle assumption wherein equiangular steps on a flat screen are assumed to correspond to equidistant points on the screen and on the stimulus itself. This is an approximation that works well for small centrally displayed stimuli that are typically 5-10 degrees of visual angle. Since the dome is hemispherical, equidistant points on the dome subtend equal angles for the viewer. Furthermore, we find that conventional 9-point calibration at the central +/-20° gives good calibration accuracy within 1-2 degrees that is well within the size of objects in our naturalistic stimuli. Thus, it is possible to both enforce central fixation and also track free-viewing eye movements in the dome environment.

### 2.4. fMRI Experiments

#### 2.4.1. Subject

An adult male Rhesus macaque (*macaca mulatta;* weight 10 kg; age 6 years) participated in the fMRI experiments. The subject was surgically implanted with a fiberglass headpost used to immobilize the head during experiments. The subject was acclimated to all aspects of the experimental procedure over a period of months, prior to data collection. All procedures were approved by the Animal Care and Use Committee of the U.S. National Institutes of Health (National Institute of Mental Health) and followed U.S. National Institutes of Health guidelines.

#### 2.4.2. Visual stimuli

##### 2.4.2.1. Checkerboard stimulus

The checkboard stimulus was generated via Python script, and consisted of a high contrast radial checkerboard, containing black and white tiles that alternated at 5° radial increments and 5° polar increments. The contrast polarity of the checkboard was inverted at a rate of 20Hz. In the ‘full’ condition, the checkerboard covered the full screen, corresponding to a horizontal field of view of >170°. In the ‘center’ condition, a mask (mid-gray) was overlaid such that only a circular region of the checkerboard subtending 20° visual angle was visible. The conditions were presented in continuous blocks of 40 seconds duration, with 3 repetitions of each condition (including ‘blank’) during each 6-minute EPI. A central fixation marker was presented throughout all conditions and consisted of a white crosshair on a mid-gray disk that subtended 2°. The subject was trained to maintain fixation within a 5° diameter window of the central fixation marker to earn liquid reward, and during fMRI scans liquid reward was given every 4 seconds provided the subject maintained the fixation requirement.

##### 2.4.2.2. Naturalistic first-person point of view video

In the naturalistic viewing experiment, the subject freely viewed movies while receiving liquid reward at regular intervals (every 4s) for keeping their eyes open. No fixation marker was presented, except at the start of each run, to check for possible drift of the eye tracker. Each experimental run consisted of 8 blocks of 40 seconds each, with 2 repetitions of each of 4 conditions: blank, original, center, and scrambled. The ‘original’ condition consisted of 10 movie clips of 4 seconds duration each, concatenated and presented continuously. Each movie clip consisted of 180° fisheye frames (2160 x 2160 pixels at 30 frames per second) of either real-world scenes (Lucid Camera) or virtual-reality renders (Blender). In all clips, the camera moved forward through the scene at a natural pace (human walking speed for real-world clips, or 2 m/s for virtual clips). In the ‘center’ condition, the same movie clips used in the ‘original’ condition were scaled down so that the entire fisheye frame (180° scene) covered a smaller central circular area of the dome surface equal to 20° retinal subtense. The ‘center’ condition therefore contained the same complete scene information and relative motion patterns as the ‘original’ but was no longer immersive and yielded no visual stimulation in the periphery beyond 10° eccentricity (consistent with many typical visual neuroscience experiments).

#### 2.4.3. Data acquisition

For fMRI experiments, the subject was seated in the sphinx position in an MR-compatible monkey chair (Rogue Research Inc., Canada), with its head stabilized by a custom fiberglass headpost clamp (Section on Instrumentation, NIMH). A custom 10-channel radiofrequency receive coil array embedded in a 3D-printed casing (tailored to the subject’s head and implant geometry) was positioned around the head. Liquid reward (apple juice) was delivered to the subject’s mouth via a computer-controlled pressurized delivery system (Mitz, 2005). The animal received reward at regular 4 second intervals throughout each run, provided their eyes remained open. Eye position was monitored via video eye tracking using an MR-compatible infrared camera (Model 12M, MRC Systems GmbH, Germany), that sent video signal to the eye tracker software (ViewPoint, Arrington Research, Az). Analog signals were routed via a data acquisition card (DataPixx 2, VPixx Technologies, Canada) and recorded in Matlab. Visual stimuli were projected from an Epson PowerLite projector in the configuration described previously (Section 2.1.1.2), and experimental timing was controlled by Matlab and synchronized to volume acquisitions by the scanner.

Echo-planar imaging (EPI) sequences were acquired at 1.2mm isotropic voxel resolution (TR = 2s; TE = 18ms, GRAPPA acceleration factor = 2). The EPI phase-encoding direction was set as left to right to minimize image distortions caused by occasional body motion. Each EPI run began with 4 dummy TRs, which were discarded from analysis. Monocrystalline iron oxide nanoparticles (MION) manufactured by the Chemistry and Synthesis Center (NHLBI) was injected intravenously to the saphenous vein immediately prior to fMRI experiments (30mg FE/mL; 9 mg/kg) to enhance MR-contrast. Anatomical T1 and T2-weighted MRI volumes were acquired in a separate scan session, during which the subject was sedated under ketamine anesthesia (15 mg/kg, intramuscular injection) and dexmedetomidine (0.01 mg/kg intramuscular), with glycopyrrolate (0.01 mg/kg, I.M.) administered to reduce excess salivation caused by ketamine. The two functional MRI experiments were run in separate sessions on different days, and the functional data were aligned to the high-res anatomical scans from the anesthetized session.

#### 2.4.4. Data analysis

MRI data were first converted from DICOM to NIFTI format using dcm2nii (Li et al., 2016) and reoriented using FSL (fslorient) to match the volume to the subject’s real-world head orientation (i.e. ‘sphinx’ position). Susceptibility distortion of the EPI was corrected using FSL’s TOPUP correction (Andersson et al., 2003). All subsequent data pre-processing was performed using AFNI (Cox, 1996; Cox & Hyde, 1997), which included motion correction, slice timing correction, up-sampling (to 1mm isotropic voxel dimensions) and finally GLM analysis (Taylor et al., 2024). A motion threshold of 0.4 mm was used to censor TRs from the regression model. For approximate anatomical identification of brain areas, the D99 macaque atlas (Reveley et al., 2016; Saleem & Logothetis, 2012) was aligned to the subject’s native anatomical using AFNI’s @AnimalWarper and the NMT 2.0 template (Jung et al., 2021).

## 3. Results

### 3.1. Whole brain response to immersive viewing

To demonstrate the utility of the immersive display for fMRI, we first asked which parts of the primate brain are preferentially activated during visual stimulation of the far periphery. To address this question, we collected fMRI data from an awake monkey during full-field (∼180° horizontal FOV) and central field (20° FOV) presentation of a flashing checkerboard stimulus, while the animal maintained central fixation. Figure 5A illustrates the contrasting hemodynamic responses in occipital cortex to these two conditions. While central stimulation elicited its strongest activation in lateral portions of the visual cortex, full-field stimulation yielded a broader response that included strong activation of medial portions of the visual cortex, especially around the calcarine sulcus on the medial surface.

**Figure 5.**
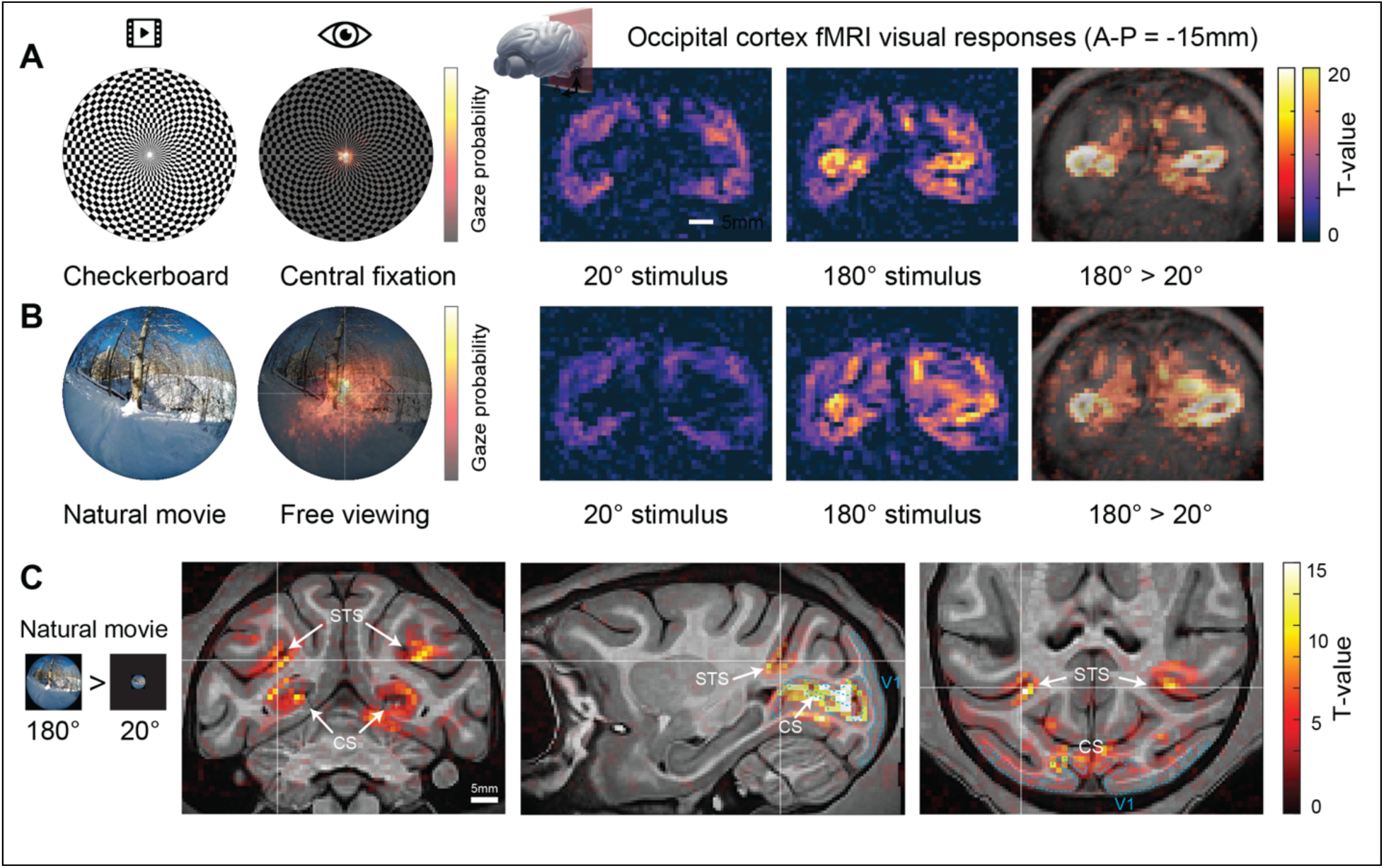
Comparison of stimulus-evoked brain responses under controlled fixation (A) and natural viewing conditions (B). (A) Checkerboard stimulus paradigm with central fixation, showing the full-field stimulus and corresponding gaze probability distribution, which is tightly concentrated around the fovea. (B) Natural movie stimulus under free viewing, with broader gaze probability distributions reflecting exploratory eye movements. In both conditions, stimulus sequences alternate between full-field (180°), blank, and center-restricted (20°) presentations. Cortical slice panels on the right show example fMRI activation maps (T-values) across the same coronal slice (AP=-15mm) for center-restricted and full-field conditions, respectively, demonstrating stronger and more widespread activation for full-field stimulation and more localized responses for central stimulation. Comparisons across conditions reveal differences in spatial extent and magnitude of activation, with free viewing of natural movies producing distributed but behaviorally modulated responses, whereas controlled fixation with checkerboard stimuli yields more stereotyped, centrally weighted activation patterns. Color bars indicate T-values, and the inset anatomical overlays (rightmost column) provide spatial reference for activation localization. (C) Detail of visual cortex stimulation evoked by full-field fisheye natural movie versus a center-restricted version. Coronal (left), sagittal (middle), and axial (right) slices show fMRI activation maps (warm colors; T-values) overlaid on anatomical images during presentation of wide-field natural movie stimuli. Prominent activation is observed in the fundus of the superior temporal sulcus (STS) and along the calcarine sulcus (CS), consistent across views (white labels and arrows). Boundaries of primary visual cortex (V1) are shown in dashed cyan based on atlas registration.

Having established a mapping of the retinotopic representation of the far peripheral visual field in visual cortex, we next asked how these areas respond under more naturalistic viewing conditions. To this end we presented movies from first person perspective of self-motion through real world scenes, while the subject viewed the stimulus freely. Correspondingly, video eye tracking revealed a broader distribution of eye gaze positions during this experiment (Figure 5B), although fixations were still biased towards the center of the display. We again contrasted responses to full-field presentation (180°) versus traditional central field stimulation (20°). The patterns of activation to each condition with the naturalistic movies were similar to those elicited by the checkboards used in the previous experiment. Specifically, medial regions of the occipital cortex, including the calcarine sulcus, responded more strongly to full-field stimulation than to central stimulation.

Activation under immersive conditions extended anteriorly from occipital cortex (Figure 5C), and beyond primary visual cortex into extrastriate areas, suggesting recruitment of larger-scale networks during naturalistic viewing of vection (i.e. simulated first-person perspective locomotion). This effect was further evident along the superior temporal sulcus (STS) and caudal visual regions (Figure 5C), which includes regions previously implicated in the visual encoding of optic flow. The spatial distribution of responses indicates engagement of motion-sensitive and higher-order visual processing regions, which are less prominently activated under conventional small-field paradigms.

## 4. Discussion

### 4.1. Summary

In this work, we introduced and validated dome visual stimulation frameworks, that enables the study of immersive naturalistic vision, under controlled experimental conditions. Building on hemispherical projection approaches (P. D. Bourke, 2005) and adapting them for electrophysiology and MRI-compatible use, we developed a system that combines wide-field projection (∼180°), geometrically accurate stimulus rendering, and calibrated pre-warping to preserve real-world metric relationships. Our pipeline integrates advanced stimulus generation (Unity/Blender-based fisheye rendering), computer-vision-based distortion correction, and MRI-compatible optical and eye-tracking solutions, thereby overcoming constraints associated with space, projection geometry, and compatibility within fMRI and electrophysiology environments.

### 4.2. Neural responses in the dome

The neural mechanisms of peripheral visual processing have historically been understudied, in part, due to methodological constraints. We therefore demonstrated of the utility of an immersive dome system by mapping fMRI responses of the primate brain to visual stimulation of the far peripheral visual field, up to 90° eccentricity. We observed that full-field visual stimulation elicited stronger fMRI responses in medial regions of occipital cortex compared to matched small-field stimuli, typical of traditional visual experiments. This result is consistent with evidence for retinotopic organization of primate visual cortex from human functional imaging (Mikellidou et al., 2017). Previous studies in macaque have identified visually-responsive regions along the calcarine sulcus on the medial surface of the cortex as corresponding to more peripheral regions of the visual field (Arcaro et al., 2022). Furthermore, the robust activation observed along the superior temporal sulcus (STS) and caudal visual regions under immersive stimulation (Fig. 5) highlights the engagement of higher-order visual processing pathways, which are often underrepresented in conventional paradigms. Together, these results demonstrate that increasing the spatial extent and ecological validity of visual input significantly alters the magnitude and distribution of brain-wide activity.

Given the retinotopic basis of this result, one might predict that this pattern of activation would change substantially under more naturalistic conditions during which the subject is free to move the eyes, and where the stimulus motion drives attention. However, the results from our second experiment using naturalistic movies revealed a remarkably similar map of preferences for full-field over reduced field stimuli. This is likely due to the extreme magnitude of difference in visual field stimulation between conditions (180° vs 20°), relative to the magnitude of differences in eye movements between experiments. We observed a central bias in gaze fixation distributions during free-viewing, despite the higher overall variability, which may be driven by the visually salient focus of expansion for optic flow typically occurring near the center of the display (Knöll et al., 2018). These findings are consistent with prior work showing that naturalistic, dynamic stimuli engage distributed cortical networks and enhance functional correlations across brain regions (S. H. Park et al., 2017; Russ & Leopold, 2015a). However, that work used traditional visual display methods that stimulated only the central <20° of the visual field and contained no first-person self-motion.

### 4.3. Novel research directions afforded by dome systems

A key conceptual advance of this work is the incorporation of naturalistic visual immersion into experimental neuroscience. The subjective feeling of immersion arises not only from wide spatial coverage but also from integration of central and peripheral vision. It is also closely connected with behavioral actions such as orienting and moving through the environment. By carefully creating visual projections that accurately preserve the scene geometry, such first-person actions provide a compelling perception of interacting with the environment. The neural mechanisms surrounding these core features of visual cognition are surprisingly unexplored and well suited for study in the dome.

One area of opportunity relates to the metric distances and sizes of objects within a scene. In the real world, objects have physical sizes, whose subtense on the retina changes as a function of distance. Recent work demonstrates that object-related areas in the inferior temporal cortex of the macaque encode metric features of objects and scenes (Khandhadia et al., 2023b; Vaziri et al., 2014b). Emulating self-motion through the environment, thus mimicking a constant source of visual input for humans and other animals, the real-world geometry of the scene is emphasized. For example, the sequential perspective on 3D objects gained by advancing through a scene disambiguates their relative positions, sizes and to some extent their volumetric shapes. Motion parallax is particularly strong in the periphery (Kim et al., 2016) and cannot be adequately reproduced using small, centrally presented stimuli. As most scene elements are fixed in their locations, the dynamic retinal stimulation consequent to self-directed actions can be interpreted relative to an unchanging backdrop. Relatedly, the continuity of time provides a context for interpreting self-directed and externally originating events. Recent work demonstrates the importance of temporal continuity in structuring the responses of neurons during natural modes of visual behavior in the macaque (Russ et al., 2023; Russ & Leopold, 2015b).

In rodent studies, immersive environments have been successfully used to study navigation and sensorimotor integration in rodents (Hölscher et al., 2005; Katayama et al., n.d.). The adoption of visual immersion has been less for non-human primate research. Although there is a push towards social housing and freely behaving primates with chronic implants (Testard et al., 2023), such designs lack parametric modulation that is possible using immersive visual displays. More recently, ultra-wide-field neuroimaging in humans has shown that immersive scene presentation recruits extended visual cortex beyond traditional retinotopic boundaries (J. Park et al., 2024). Our results extend these findings to non-human primates and demonstrate that immersive, full-field stimulation engages parts of the visual cortex that may more closely resembles real-world vision. Importantly, an embodied representation of one’s environment based on motion signals may convey metric information about real-world structure that is not captured by static image features alone, highlighting the limitations of image-based approaches to studying vision.

### 4.4. Limitations

The dome display systems can deliver dynamic full-field color visual stimulation, but there are some trade-offs that could pose limitations for certain experimental scenarios. A fundamental limitation of using a single fixed projector to illuminate a hemispheric dome is that the pixel resolution and luminance will naturally decrease with increasing eccentricity from the optical axis. For our purposes of naturalistic full-field stimulation, these effects were not deemed perceptually salient to human observers, partially aided by the fact that visual acuity is lower in the periphery. Correction of luminance variation across the display is possible by taking photometric measurement across the dome surface to calculate a color look-up table to adjust output at each pixel location. This level of calibration may be important for certain experiments.

Given the importance of visual depth perception to naturalistic immersive vision, an obvious beneficial addition to dome display systems would be the addition of stereoscopic depth cues through binocular stimulation. We initially designed both dome systems to present stereoscopic 3D stimuli via passive polarization filter systems (Boothroyd, 2010). In the context of neuroscience experiments, passive stereoscopic presentation methods are often preferred to keep active electronics away from neural recording electrodes or the magnetic field. However, we found that despite the careful use of appropriate commercial 3D screen surface treatments (Zhang et al., 2012), internal reflections (front-surface dome) and refraction (rear-surface dome) negatively impacted light polarization, resulting in cross-talk and ghosting (Woods, 2012) that diminishes the 3D percept. Alternative approaches to passive binocular presentation that do not rely on polarization, such as color anaglyph or interference filters, may produce more acceptable stereoscopic results, albeit at the expense of accurate depiction of color.

Finally, although naturalistic stimuli provide greater ecological validity, they reduce experimental control relative to traditional paradigms and make it more challenging to isolate the specific stimulus features driving observed neural responses. This is where the use of computer generated (virtual reality) scenes holds tremendous advantage over real world video footage. Precise parametric manipulation of nearly any variable is possible in the virtual environment, which presents both unparalleled opportunities but also new challenges to researchers seeking the most scientifically important and behaviorally relevant aspects of natural vision to study.

### 4.5. Future Directions and Conclusion

Beyond basic sensory processing, the dome framework provides a powerful platform for investigating ecologically relevant visual behaviors. Accurate perception of object size, distance, and motion is critical for navigation, foraging, and social interaction, yet these aspects are often simplified or ignored in traditional experiments. Compared to head-mounted display (HMD) systems, which are limited in field of view and can introduce perceptual artifacts (Maeda et al., 2024; Xiao et al., 2025b), dome-based displays offer a more naturalistic and scalable solution, particularly for non-human primates with different anatomical constraints (Bush & Miles, 1996). At the same time, dome systems retain the experimental control necessary for systematic manipulation of stimuli, allowing researchers to bridge the gap between controlled laboratory studies and real-world visual experience. Future work can also leverage this framework to investigate higher-order processes such as scene understanding, multisensory integration, and behaviorally relevant perception, ultimately advancing our understanding of how the brain operates in real-world environments.

In striving for increased ethological validity while retaining the level of experimental control required for ease of interpretation of results, immersive visual stimulation can cover a sweet spot. By incorporating visual immersion, peripheral stimulation, and self-motion cues into experimental design, dome-based systems provide a powerful approach for studying natural vision.

## Acknowledgements

We are grateful for the MRI assistance from Charles Zhu and Frank Ye from the Neurophysiology Imaging Facility, 3D design and printing from Camille Rood in the NIMH Section on Instrumentation, help with fMRI analysis pipelines from Paul Taylor and Daniel Glen from the NIMH Scientific and Statistical Computing Core, and provision of MION contrast agent from Haitao Wu and Rolf Swenson from the NHLBI Chemistry and Synthesis Center. This research was supported by the Intramural Research Program of the National Institutes of Health (NIH) (ZIAMH002838 and ZICMH002899). The contributions of the NIH authors are considered Works of the United States Government. The findings and conclusions presented in this paper are those of the authors and do not necessarily reflect the views of the NIH or the U.S. Department of Health and Human Services.

## Contributions

**Harish Katti**: Conceptualization, Investigation, Methodology, Project administration, Software, Visualization, Writing **Aidan P. Murphy**: Conceptualization, Data curation, Formal analysis, Investigation, Methodology, Validation, Visualization, Writing **Michael Helde**: Data curation, Formal analysis, Validation, Software, Writing **Harshawardhan Deshpande**: Data curation, Formal analysis, Investigation, Validation, Visualization, Writing **Tyler J. Lee**: Data curation, Resources **Rekeya Knight**: Data curation, Resources **Carly Gregg**: Resources **Cristina Solinas**: Resources **Katherine Cameron**: Resources **Daryl Bandi**: Resources **George Dold**: Resources **David A. Leopold**: Conceptualization, Funding acquisition, Investigation, Methodology, Project administration, Resources, Software, Supervision, Validation, Visualization, Writing

