## Supplementary Materials for "Immersive display systems for simulating natural vision"

### Supplementary Material

1. Setting up a 180° fisheye camera in Blender
2. Setting up a 180° fisheye camera in Unity
3. Front-surface hemispheric dome screen calibration
4. A Comparison of Software Tools

#### 1. Setting up a 180° fisheye camera in Blender

Every parameter of a Blender (www.blender.org) scene can be controlled via the graphical user interface or programmatically via Python script:

Manual configuration via graphical user interface

For 180° fish-eye renders in Blender (v5.0), the 'Cycles' render engine should first be selected. With a camera object selected, in the camera 'Data' tab of the Properties menu, the lens type should be set to 'panoramic' and 'panorama type' should be set to 'fisheye equidistant' (Figure S1), which then allows adjustment of 'field of view' (°). The aspect ratio of the output frame images should be set at 1:1, with the pixel values matching (or exceeding) the vertical resolution of the projector. The rendered output will appear as a fisheye image within a circular aperture and black pixels in the corners of the image.

Configuration via Pyhton script (FisheyeSetup.py)

```
import math
import bpy

bpy.data.scenes["Scene"].render.engine = 'CYCLES'
bpy.data.scenes["Scene"].render.resolution_x = 2160
bpy.data.scenes["Scene"].render.resolution_y = 2160
bpy.data.cameras["Camera"].type = 'PANO'
bpy.data.cameras["Camera"].panorama_type = 'FISHEYE_EQUIDISTANT'
bpy.data.cameras["Camera"].fisheye_fov = math.radians(180)
```

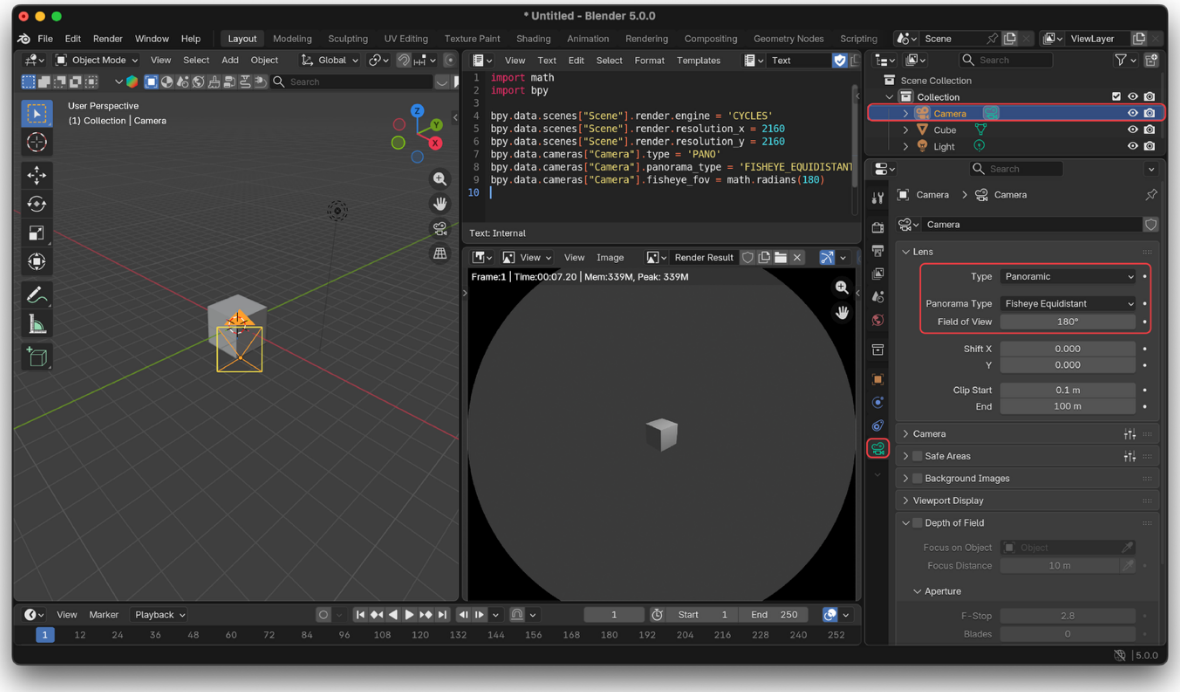


**Figure S1.** A screenshot of the Blender (v.5.0.0) UI with default scene loaded. By selecting the 'camera' object (highlighted orange in the '3D Viewport' window on the left, and highlighted in red in the 'Outliner' window in the top right), you can manually change the camera's lens parameters, as shown in the 'Properties' window on the right. Alternatively, all aspects of Blender can be controlled programmatically via Python scripts, as seen here loaded into a 'Text Editor' window (middle column, top row). When the scene is rendered, the output frames are 180° fisheye projections, which appear as a circular image (middle column, bottom row).

#### 2. Setting up a 180° fisheye camera in Unity

For generating 180° fisheye output from the Unity (www.unity.com) game engine, we provide the C# script 'SimpleFisheyeCamera.cs'. This is an implementation of a cubemapped fisheye projection to obtain 180° full field of view coverage of the scene (see Figure S2). Attach SimpleFisheyeCamera.cs to the main camera in your Unity project, and assign the CustomFisheye.mat shader to this camera.

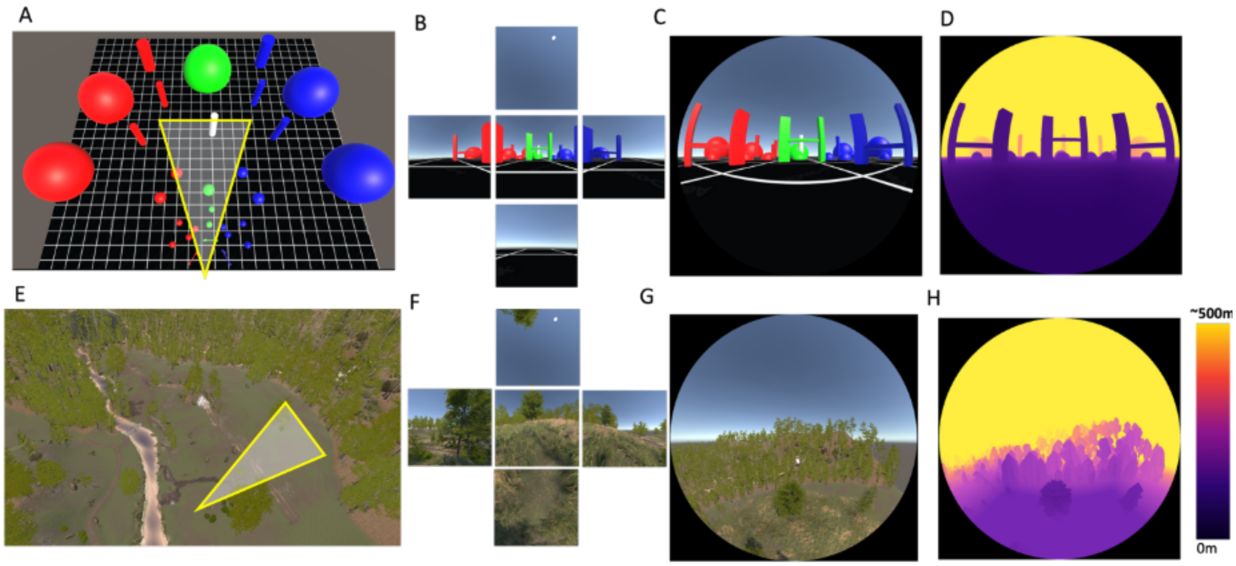


**Figure S2. Cubemap-based fisheye rendering and depth recovery in synthetic 3D environments.** (A) Bird's-eye view of an abstract 3D scene in Unity®, consisting of colored geometric objects arranged on a ground plane. The yellow triangle indicates the viewing frustum corresponding to the virtual fisheye camera. (B) Five orthogonal camera views used for cubemap generation (front, left, right, top, and bottom), which together sample the full visual environment surrounding the observer. (C) Final fisheye image generated by remapping the cubemap into an equidistant hemispheric projection suitable for dome display and immersive visual stimulation. (D) Ground-truth depth map recovered directly from the rendering pipeline and scene geometry associated with the fisheye image; color shades depict depth from observer. (E–H) The same rendering pipeline applied to a naturalistic outdoor 3D environment in Unity®. (E) Bird's-eye view showing the fisheye camera viewing frustum within the scene. (F) Corresponding cubemap camera views sampling the environment. (G) Generated fisheye rendering of the naturalistic scene. (H) Ground-truth depth map corresponding to the fisheye image, illustrating recovery of scene depth structure from the rendering process, color shades depict depth from observer. Together, these panels illustrate the cubemap-to-fisheye rendering framework used for immersive dome stimulus generation and associated extraction of metric depth information from synthetic virtual environments. A similar process was used to generate Figure 1 e, f, g, using Blender®.

### 3. Front-surface hemispheric dome screen calibration

The provided MATLAB script '`dome_projection_calibration_and_warping.m`' calibrates the geometric distortion produced when visual stimuli are projected from a flat projector image onto a curved dome, and then transfers the learned correction into a Psychtoolbox (http://psychtoolbox.org/) calibration file. The script first creates a polar set of calibration locations in projector coordinates using concentric eccentricity rings and radial directions. Each calibration point is displayed individually with Psychtoolbox as a red dot, and a webcam records where that projected point actually appears on the dome. A second image is acquired after the dot is removed, and background subtraction is used to identify the dot location in camera coordinates. These measured locations are converted from camera pixels into the same coordinate system as the projector image. The resulting pairs of intended locations, stored in refPts, and measured projected locations, stored in domePts, define the geometric distortion of the complete projector and dome system.

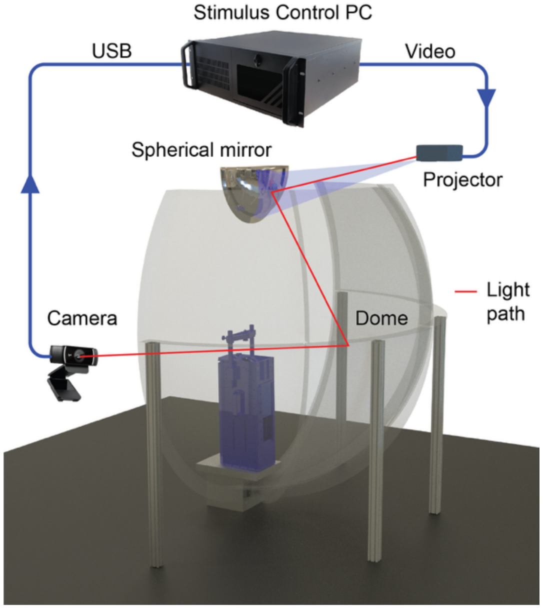


The script then learns both a forward mapping from intended screen coordinates to measured dome coordinates and an inverse mapping from measured dome coordinates back to the projector coordinates required to produce the intended location. The mapping can be estimated either with robust second-order polynomial regression or with thin plate splines. The learned calibration, calibration points, screen dimensions, and dome geometry parameters are stored in calibtex.mat so that the physical calibration procedure does not need to be repeated for subsequent experiments.

Finally, the script loads an existing Psychtoolbox calibration file, calibtestfilex.mat, rescales its calibration-dot coordinates to the 3840 by 2160 projector resolution, applies the learned inverse mapping to each dot, replaces XCALIBDOTS and YCALIBDOTS with the corrected positions, and saves the resulting standard calibration structure as calibtestfilex_warped.mat. This allows the empirically measured dome correction to be used by the existing Psychtoolbox stimulus presentation pipeline.

**Key variables**

**`screenWidth and screenHeight`**

The native projector resolution in pixels. In this script they are 3840 and 2160. These values define the coordinate system in which calibration points and corrected Psychtoolbox locations are represented.

**`ringAnglesDeg`**

The eccentricities used to generate the concentric calibration rings. The current setting, 5:5:90, samples locations from 5 to 90 degrees in 5 degree increments. The corresponding radius in projector pixels is calculated as (screenHeight/2) * sind(angle).

`lineAnglesDeg`
The polar directions used to sample the calibration grid. The current setting, 0:10:350, samples radial directions every 10 degrees around the display.

`screenPts`
An N by 2 array containing the intended calibration locations in projector coordinates. Each row contains an x and y location at which a calibration dot is displayed.

`domePtsCam`
An N by 2 array containing the detected locations of the projected calibration dots in webcam coordinates. These are the raw measurements of where the projector actually places each requested point on the dome.

`domeCenterX and domeCenterY`
The location of the dome center in the webcam image. These values are used to recenter the webcam measurements before converting them into projector-scale coordinates.

`domeRadiusPixels`
The effective dome radius measured in webcam pixels. Together with domeCenterX and domeCenterY, this variable determines the conversion between webcam coordinates and the projector coordinate system.

`domePts`
The measured dome locations after the webcam coordinates have been recentered and rescaled into projector-coordinate units. These coordinates can therefore be directly compared with screenPts.

`refPts`
The desired or reference coordinates for calibration. In this script refPts is assigned directly from screenPts. The correspondence refPts(i,:) to domePts(i,:) describes how the projection system displaced calibration point i.

`mappingMethod`
Selects the mathematical model used to represent the distortion. It can be set to 'poly' for robust second-order polynomial fitting or 'tps' for thin plate spline fitting.

`calib.fx and calib.fy`
For polynomial calibration, these models describe the forward mapping from desired screen coordinates to the x and y coordinates measured on the dome.

`calib.gx and calib.gy`
For polynomial calibration, these models describe the inverse mapping from measured dome coordinates to the projector coordinates required to obtain the desired location. These inverse models are the mappings used to modify the Psychtoolbox calibration dots.

`calib.tps_fx and calib.tps_fy`
Thin plate spline versions of the forward screen-to-dome mapping.

`calib.tps_gx and calib.tps_gy`
Thin plate spline versions of the inverse dome-to-screen mapping. These are used when mappingMethod is set to 'tps'.

`calib`
A structure containing the learned mapping together with the calibration correspondences and geometric parameters. It is saved to calibtex.mat and can be loaded in later sessions by setting performCalibration to false.

`performCalibration`
Controls whether a new physical calibration is performed. When true, calibration points are projected and measured with the webcam and a new model is learned. When false, the previously saved calibration is loaded from calibtex.mat.

`calibFile`
The filename used to store the learned calibration. It is currently set to calibtex.mat.

`ptbCalibFile`
The input Psychtoolbox calibration file. It is currently calibtestfilex.mat and is expected to contain the structure scal with the fields XCALIBDOTS and YCALIBDOTS.

`xy`
The Psychtoolbox calibration-dot coordinates after conversion from the coordinate convention in calibtestfilex.mat to the 3840 by 2160 projector coordinate system. These are the locations to which the learned inverse dome correction is applied.

`scalxpred and scalypred`
The corrected x and y projector positions predicted by the inverse calibration mapping for the Psychtoolbox calibration dots.

`c.scal.XCALIBDOTS and c.scal.YCALIBDOTS`
The standard Psychtoolbox calibration-dot fields. The script replaces their original values with scalxpred and scalypred before saving the new calibration file.

`calibtestfilex_warped.mat`
The final Psychtoolbox calibration file containing the corrected calibration-dot locations. This file incorporates the empirically learned projector-to-dome distortion correction and can be used by the subsequent Psychtoolbox presentation pipeline.

`prewarp_frontprojection_dome.m`
(Needs psych toolbox installed on Matlab)

PsychToolbox (PTB) based routine that creates a Screen object that has calibration data in 'PTB_calibfile_for_frontprojection_dome_prewarp.mat' as an attribute. The source videos and images are then undistorted/ pre-warped to be suitable for projection onto front projection dome. The resolution of front projection dome should be set to 1920x1080 for this output to look correct.

#### 4. A Comparison of Software Tools

| Experimental / Technical Area | Python Scripts | Psychtoolbox (MATLAB) | MonkeyLogic | Unity | Blender |
|---|---|---|---|---|---|
| Primary Role | analysis, preprocessing, machine learning, and stimulus generation | controlled stimulus presentation and records behaviour | controlled stimulus presentation and records behaviour | immersive real-time rendering and VR | Supports offline asset and animation creation |
| Natural and Synthetic Stimuli | moderate support for naturalism | less support for naturalism | less support for complex graphics | Supported | Supported |
| Static and Dynamic Stimuli | Supported | Supported | Supported | Supported | Supported |
| Real-time Interactive VR | Less support | Less support | Not supported | Supported | No support |
| Self-motion, Navigation, and Optic Flow | Supported | Some support | Less support | Supported with real-time immersive simulation | Less support |
| Physics, Environmental Interaction, and Procedural Worlds | Less support | No support | No support | Supported in real time | Supported for offline rendering |
| Stimulus quality | Moderate, depends on library used | High quality for parameterized stimuli, e.g.; Gabors | Good quality for simple parametric stimuli | Moderate/high quality | High quality |
| Timing Precision, Trial Logic, and Behavioral Control | Less support for deterministic timing | Supported | Supported | Less support | No native support |
| Eye Tracking | Supported, needs programming | Supported natively | Supported natively | Less support, needs programming | No native support |
| Fisheye, Cubemap | Supported with custom programming | Supported with image warping | Less support | Supported natively and efficiently | Supported mainly for offline rendering |
| Typical Role in a Dome Experiment Pipeline | Analysis backend and computational tools | Precise visual presentation, behaviour control | Behavioral experiment controller | Immersive VR and rendering engine | Asset and environment production tool |

**Table S1:** Summary of the typical roles of Python scripts, Psychtoolbox, MonkeyLogic, Unity, and Blender in immersive dome-based neuroscience and vision experiments.

These tools are often used together rather than as isolated systems, with each platform contributing different strengths to stimulus generation, behavioral control, rendering, analysis, and asset creation. The comparison focuses especially on the types of workflows explored in the fisheye and dome-rendering projects discussed earlier, including naturalistic environments, optic flow, self-motion, and social-agent stimuli.

- **Python** is flexible and is especially useful for generating and processing natural movies, computing optic flow and depth-related regressors, etc. While Python can generate stimuli directly through OpenGL or PsychoPy-based systems, it is less commonly used for tightly timed online presentation compared to Psychtoolbox or Unity.

- **Psychtoolbox**, is widely used for presenting carefully controlled visual stimuli such as drifting gratings, looming stimuli, sparse noise, random-dot fields, retinotopy patterns, and controlled optic-flow displays. In dome systems, Psychtoolbox is often combined with custom fisheye warping pipelines so that visual stimuli are projected correctly onto hemispherical displays. Its major strength is deterministic frame timing and reliable synchronization with electrophysiology systems, photodiodes, and eye trackers. However, it is less suited for large immersive virtual worlds or complex interactive environments.
- **MonkeyLogic** is primarily a behavioral control framework rather than a rendering engine. It excels at managing trial structures, fixation requirements, reward delivery, eye tracking, state-machine logic, and experimental sequencing, especially in primate experiments. In immersive dome experiments, MonkeyLogic is often paired with either Psychtoolbox or Unity. In such setups, MonkeyLogic controls task logic and behavioral timing while Unity or Psychtoolbox generates the visual environment. MonkeyLogic itself has less support for complex rendering, but it provides strong support for behavioral experiment control and integration with neural recording hardware.
- **Unity** can serve as a primary engine for immersive real-time dome rendering and interactive virtual reality experiments. It is particularly well suited for naturalistic environments, self-motion experiments, navigation tasks, optic-flow paradigms, and socially relevant animated agents. The fisheye and cubemap rendering pipelines discussed earlier fit naturally within Unity because the engine already supports real-time 3D rendering, cubemap generation, physics simulation, dynamic lighting, animated avatars, procedural environments, and camera trajectories. Unity is especially strong for experiments involving forward motion, motion parallax, navigation through realistic environments, or social-agent interactions such as approaching macaques or animated human avatars. Compared to Psychtoolbox, Unity sacrifices some deterministic timing precision but gains much greater flexibility for immersive real-time virtual environments.
- **Blender** occupies a different role from the other systems. It is primarily an offline asset-creation and animation platform rather than an online experimental engine. Blender is particularly useful for constructing high-quality 3D environments, rigging and animating characters, generating cinematic sequences, creating procedural geometry, and preparing models for export into Unity. Blender supports offline fisheye rendering and cubemap rendering, but it is generally not suitable for closed-loop neuroscience experiments or behavioral control.

Self-motion and optic-flow experiments form a particularly important category in immersive dome systems. 3D engines such as Blender and Unity are especially strong in this area because camera motion through a real 3D environment automatically generates physically consistent optic flow, motion parallax, and depth-dependent image transformations. In contrast, Psychtoolbox typically generates optic-flow fields mathematically, which provides strong experimental control but less environmental realism. Timing precision and behavioral control remain major reasons why Psychtoolbox and MonkeyLogic continue to be widely used in neuroscience laboratories. Psychtoolbox provides highly reliable frame timing and synchronization with hardware, while MonkeyLogic provides strong support for trial sequencing, reward logic, and eye tracking. Unity can support behavioral experiments but typically requires extra efforts for deterministic timing reliability.
